# Transcriptome profiling identifies the chromatoid body as a dynamic center for RNA surveillance and transcript sorting during spermatogenesis

**DOI:** 10.64898/2026.09.02.744273

**Authors:** Lin Ma, Ammar Ahmedani, Simang Champramary, Samuli OM Laasanen, Opeyemi Olotu, Noora Kotaja

## Abstract

The chromatoid body (CB) is a hallmark of haploid male germ cells, but how it selects and regulates RNAs has remained unclear. Here we define the RNA landscape of the CB by transcriptome and small RNA profiling of isolated CBs throughout mouse round spermatid differentiation. We find that the CB selectively concentrates pachytene piRNAs, retrotransposon transcripts, specific mRNA isoforms and intron-retaining mRNAs, revealing extensive RNA sorting within this cytoplasmic germline condensate. The enrichment of transposable element transcripts together with PIWI–piRNA complexes identifies the CB as a surveillance center that may safeguard genome integrity in post-meiotic haploid cells. Unexpectedly, mRNA recruitment is not determined by predicted piRNA targeting. Instead, transcript localization is encoded by a combinatorial set of intrinsic sequence and structural features that accurately predict CB enrichment at isoform resolution. These findings establish the CB as a dynamic post-transcriptional regulatory compartment that integrates genome surveillance, RNA quality control and selective transcript sorting, uncovering general principles through which biomolecular condensates shape cell-specific transcriptomes.

## INTRODUCTION

Spermatogenesis depends on the precise temporal and spatial regulation of gene expression to ensure the continuous production of functional spermatozoa capable of fertilization and intergenerational transmission of information. In addition to the transcriptional regulation with critical role in orchestrating germ cell differentiation, post-transcriptional regulatory mechanisms play essential roles in shaping the germ cell transcriptome as spermatogenesis progresses ^1^. Post-transcriptional regulatory mechanisms become particularly critical during meiosis, a stage characterized by widespread and permissive transcriptional activity that generates a highly complex and heterogeneous RNA pool ^2,3^. In post-meiotic germ cells, this challenge is compounded by the progressive transcriptional silencing that accompanies chromatin condensation, necessitating long-term mRNA storage and tight uncoupling of transcription from translation to ensure timely protein synthesis during haploid differentiation ^4^.

A central structure implicated in post-transcriptional regulation in male germ cells is the chromatoid body (CB), a large germ cell–specific ribonucleoprotein (RNP) granule that functions as a cytoplasmic hub for RNA regulation in haploid germ cells ^5,6^. The CB begins to assemble in late pachytene spermatocytes as precursor granules, which subsequently fuse to form a single perinuclear CB shortly after meiosis in early round spermatids. This structure persists throughout round spermatid differentiation from steps 1 to 8 and disassembles upon the onset of spermatid elongation ^6^. The dynamic appearance of the CB coincides with the developmental window in which extensive post-meiotic RNA remodeling is required.

Our previous work, including our establishment of a protocol for the isolation of CBs from mouse testes, has enabled detailed molecular characterization of this unique germ granule ^7–9^. Proteomic analyses revealed that the CB is highly enriched in RNA-binding proteins and RNA-processing factors, consistent with its proposed regulatory functions. Correspondingly, transcriptomic profiling demonstrated that the CB concentrates a complex repertoire of RNAs, including mRNAs, intergenic and repeat-derived transcripts, as well as multiple classes of small non-coding RNAs (sncRNAs), most prominently PIWI-interacting RNAs (piRNAs) ^7,9^. These findings positioned the CB as a key regulatory center coordinating post-meiotic RNA fate decisions during spermiogenesis.

Among the pathways functionally associated with the CB, the piRNA pathway is particularly prominent. piRNAs are essential for safeguarding genome integrity in the fetal germline through the silencing of transposable elements ^10^, and accumulating evidence indicates that piRNAs and PIWI proteins also participate in the post-transcriptional regulation of protein-coding mRNAs in differentiating male germ cells ^11^. Notably, the CB does not appear to serve as a primary site of piRNA biogenesis ^7^. Instead, primary piRNA processing predominantly occurs earlier, in meiotic cells, at mitochondrial membranes within another germ granule termed the intermitochondrial cement (IMC) ^12–14^. During late meiosis, the dissociation of mitochondrial clusters coincides with the emergence of CB precursors. These structures are devoid of core primary piRNA processing components but become enriched in the PIWI proteins PIWIL1/MIWI and PIWIL2/MILI loaded with mature piRNAs ^7^, suggesting a developmental transition from IMC-associated piRNA production to CB-mediated piRNA effector functions in haploid germ cells.

Beyond piRNA-mediated regulation, the CB also appears to interface with broader RNA quality control and surveillance pathways, as well as mRNA storage and translational delay ^6^. Components of the nonsense-mediated decay (NMD) pathway accumulate in the CB, and we have previously demonstrated that the NMD endonuclease SMG6 is a critical CB component that contributes to shaping the transition from meiotic to post-meiotic gene expression programs ^9^. Together, existing studies support a model in which the CB functions as an integrative platform coordinating multiple post-transcriptional regulatory axes in haploid germ cells. However, the temporal dynamics of CB-associated RNA populations and the mechanisms through which distinct regulatory pathways are coordinated within the CB remain poorly understood.

In this study, we sought to provide a comprehensive characterization of CB-enriched RNAs in the mouse and to determine how CB RNA composition changes during round spermatid differentiation. By integrating RNA-seq and small RNA-seq data from isolated CBs and corresponding input round spermatid populations derived from adult testes and juvenile testes representing defined developmental stages, we delineate dynamic changes in CB-associated RNA cohorts. Our analyses provide new insights into the role of the CB in coordinating RNA surveillance and piRNA-mediated silencing during post-meiotic germ cell differentiation.

## RESULTS

### piRNAs accumulate in the CB together with PIWI proteins

Given the established role of the CB in the piRNA pathway ^7^, we first characterized the piRNA-related composition of the CB. To define the CB-associated piRNA profile, we sequenced small RNAs from adult CBs isolated by our well-established CB isolation protocol ^7^. We also sequenced the CBs and round spermatids isolated from 21 days post-partum (dpp) and 26 dpp testes (Supplementary Fig. S1A, Supplementary Table S1). These time points capture the early (21 dpp) and late (26 dpp) phases of round spermatid differentiation during the first wave of spermatogenenesis, allowing us to follow CB piRNA dynamics across differentiation. For comparison, we also enriched another type of germ granule, the IMC, using antibody against PIWIL2 ^12^ for small RNA-seq (Supplementary Fig. S1A). The mitochondria-associated IMC precedes the appearance of the CB, and it has been shown to accumulate mitochondria membrane-anchored piRNA processing machinery ^12^. Small RNA-seq of 21 dpp and 26 dpp round spermatids revealed the expected read-length distribution, with a dominant piRNA-sized population and smaller populations of other small RNAs. Biotype annotation confirmed that the round spermatid libraries contained predominantly piRNAs together with smaller fractions of miRNAs and tRNA-derived small RNAs (tsRNAs) (Fig. 1A, Supplementary Fig. S1B). In contrast, the CB samples isolated from the same developmental time points showed only the predominant piRNA peak, indicating the specific enrichment of piRNAs in the CB (Fig. 1B, Supplementary Fig. S1C). Similarly, the IMC also enriched a prominent piRNA population (Supplementary Fig. S1D) We next restricted the analysis to small RNA reads mapping to the previously annotated piRNA clusters ^15^. These reads exhibited the characteristic 5′-terminal uridine (1U) bias, a hallmark of piRNAs produced by the primary piRNA biogeneis pathway ^16,17^ (Supplementary Fig. S2A-E). The length distribution analysis showed a 24-26 nt peak for IMC-localized piRNAs, and a bit longer 29-31 nt peak for the CB-localized piRNAs (Fig. 1C). These distributions are consistent with the established association of PIWIL2/MILI with shorter piRNAs and PIWIL1/MIWI with longer piRNAs, as well as the reported localization of PIWIL2 to the IMC and early CBs, and the predominance of PIWIL1/MIWI in CBs during later steps of round spermatid differentiation ^12^ (Supplementary Fig. S3A).

**Figure 1.**
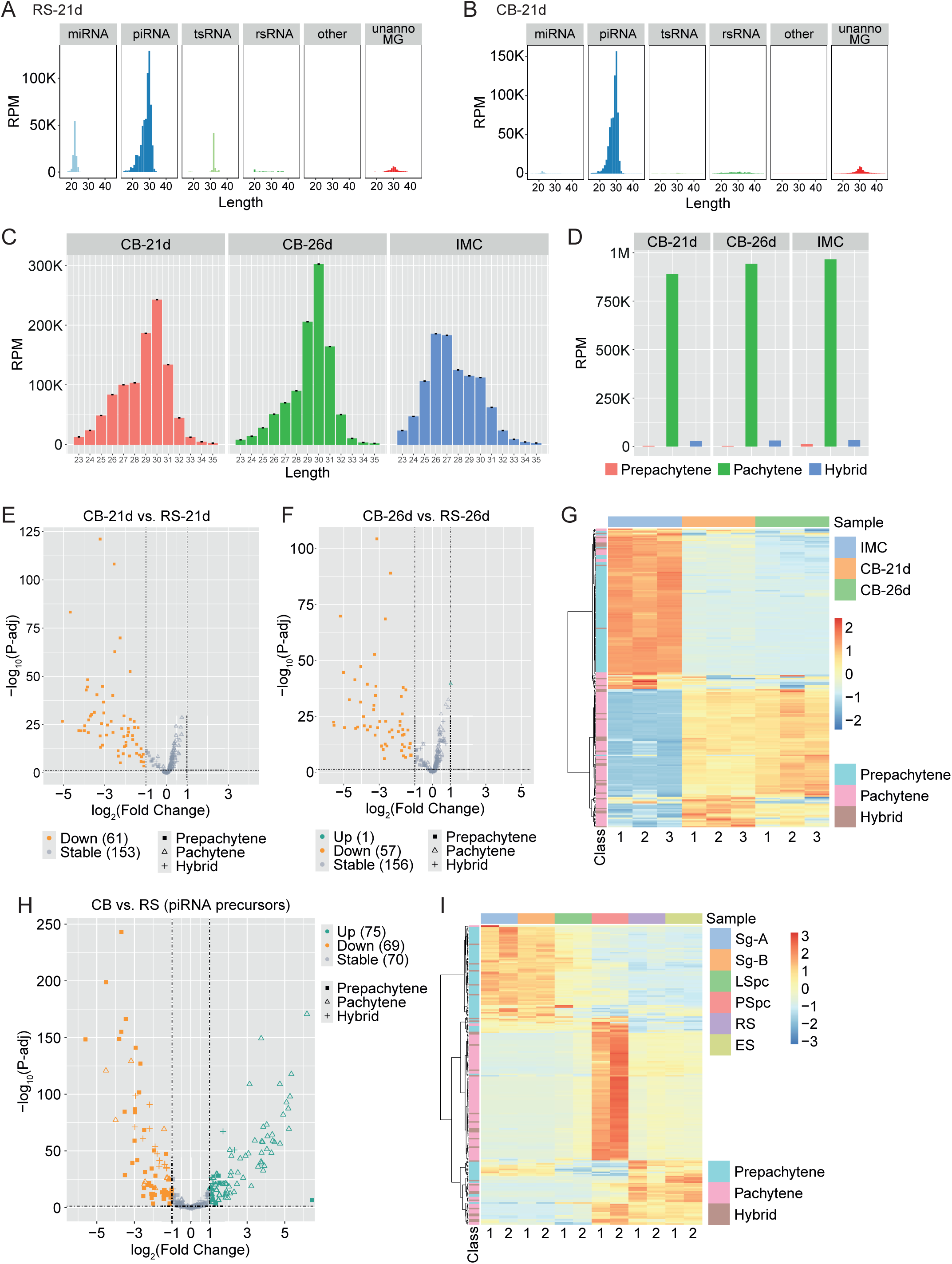
Pachytene piRNAs are enriched in the CB. (A, B) Distribution of small RNA-seq reads from round spermatid (RS)-21d (A) and CB-21d (B) between the specific small RNA subtypes. miRNA: microRNA; piRNA: PIWI-interacting RNA; tsRNA: tRNA-derived small RNA; rsRNA: ribosomal RNA-derived small RNA; unanno MG: unannotated reads matching to the genome; RS: round spermatid. (C) Length distribution of piRNAs from the CB-21d, CB-26d and IMC samples. (D) Expression levels of pre-pachytene, pachytene and hybrid piRNAs in CB-21d, CB-26d and IMC samples. (E, F) Volcano plots showing differentially expressed piRNA clusters between CB-21d and RS-21d (E) and CB-26d and RS-26d (F) (P-adj ≤ 0.05, |log₂FC| ≥ 1). (G) Heatmap showing the relative abundance of pre-pachytene, pachytene and hybrid piRNA cluster-mapping piRNAs in IMC, CB-21d and CB-26d. (H) Volcano plot showing differential expression of piRNA precursors between adult CB (batch 1) and RS (P-adj ≤ 0.05, |log₂FC| ≥ 1). (I) Heatmap showing expression patterns of piRNA precursors across spermatogenic cell types using a published dataset (GSE35005).

### CB is associated with pachytene piRNA population that remains stable during differentiation

When classified by the piRNA cluster type, piRNA reads from pachytene clusters were markedly more abundant in CBs and IMC than those derived from pre-pachytene clusters (Fig. 1D). The cluster-level differential expression analysis revealed that piRNAs originating from pre-pachytene clusters were not only present at lower overall levels but were also preferentially depleted from CBs relative to input round spermatid samples; altogether 61 and 57 clusters were downregulated in 21 dpp and 26 dpp CB compared to round spermatid samples from respective time points, and these represented almost exclusively pre-pachytene piRNA clusters (Fig. 1E,F). The cluster-level comparison of the IMC, 21 dpp CB and 26 dpp samples showed that the CB samples resembled each other with relatively higher levels of piRNAs from pachytene clusters compared to the IMC (Fig. 1G). The IMC, on the other hand, had more abundant piRNAs from pre-pachytene clusters (Fig. 1G). The direct comparison of early and late-phase CBs (26 dpp vs. 21 dpp) confirmed that the pachytene piRNA clusters are equally represented in CBs from different developmental time points, but the abundance of piRNAs from pre-pachytene clusters further decreased in the CB during differentiation (Supplementary Fig. S3B,C). Collectively, these results suggest that the CB specifically enriches pachytene piRNAs, and the CB-associated pachytene piRNA population remains quite stable during the whole course of round spermatid differentiation.

We also re-analyzed the RNA sequencing (RNA-seq) data of CBs that were isolated from round spermatids of adult mouse testes (n = 4, GSE182518) ^7,9^ to study whether, in addition to mature piRNAs, also the piRNA precursors are enriched in the CB. The differential expression analysis was performed comparing the CB samples to the negative control rabbit IgG IP samples, as well as to the CB-containing round spermatid input samples from adult testes (n = 3, GSE182518). We showed that 75 (out of 214) piRNA precursor transcripts were significantly enriched in CBs compared to round spermatids (Fig. 1H). The majority of them are precursors derived from the pachytene piRNA clusters (64 out of 75), while the pre-pachytene piRNA precursors were generally depleted from the CB (Fig. 1H). Similar enrichment was observed when compared to IgG controls, in which 40 precursors, again dominated by pachytene precursors (28), were selectively enriched in the CB (Supplementary Fig. S3D). This result is consistent with the expression pattern of piRNA precursors during spermatogenesis; pre-pachytene piRNA precursors are expressed at higher levels during earlier phases of spermatogenesis, while pachytene piRNA precursor expression peak coincides with the CB appearance (Fig. 1I). The reason for enrichment of the piRNA precursor transcripts in the CB remains unclear. Given that the CB lacks the core piRNA processing machinery, it is unlikely that theses transcripts are actively recruited to the CB for piRNA processing. Instead, considering the close developmental and functional relationship between the different types of germ granules ^13^, piRNA precursor transcripts may be carried over the CB from the IMC, a piRNA-producing compartment that precedes the CB appearance.

### CB accumulate reads from exons, introns and repeats

Next, we analysed the RNA-seq data in more detail to get novel insight into the functional role of the CB, and to identify CB-localized RNAs that are potentially targeted by piRNAs. Approximately 25% of the CB RNA-seq reads mapped to the unannotated regions of the genome (Fig. 2A), in line with our earlier observations ^7^. The identity and biological significance of these unannotated reads remain to be determined in the future studies. In the present study, we focused on annotated genomic regions and found that a considerable fraction of CB-associated reads originated from rRNA, repetitive elements, and both exonic and intronic regions of protein-coding genes (Fig. 2B). To facilitate comparison with round spermatid input samples generated by an rRNA-depleted mRNA-seq pipeline, rRNA reads were excluded from the analysis. Global RNA profiling revealed that CB-associated RNA composition differs markedly from the whole round spermatid transcriptome, with CB samples showing increased representation of repetitive elements, piRNA clusters and intronic sequences, whereas round spermatid samples were dominated by exonic reads (Fig. 2C). This finding indicates that the CB does not simply reflect the global RNA composition of round spermatids but instead selectively accumulates a distinct subset of transcripts.

**Figure 2.**
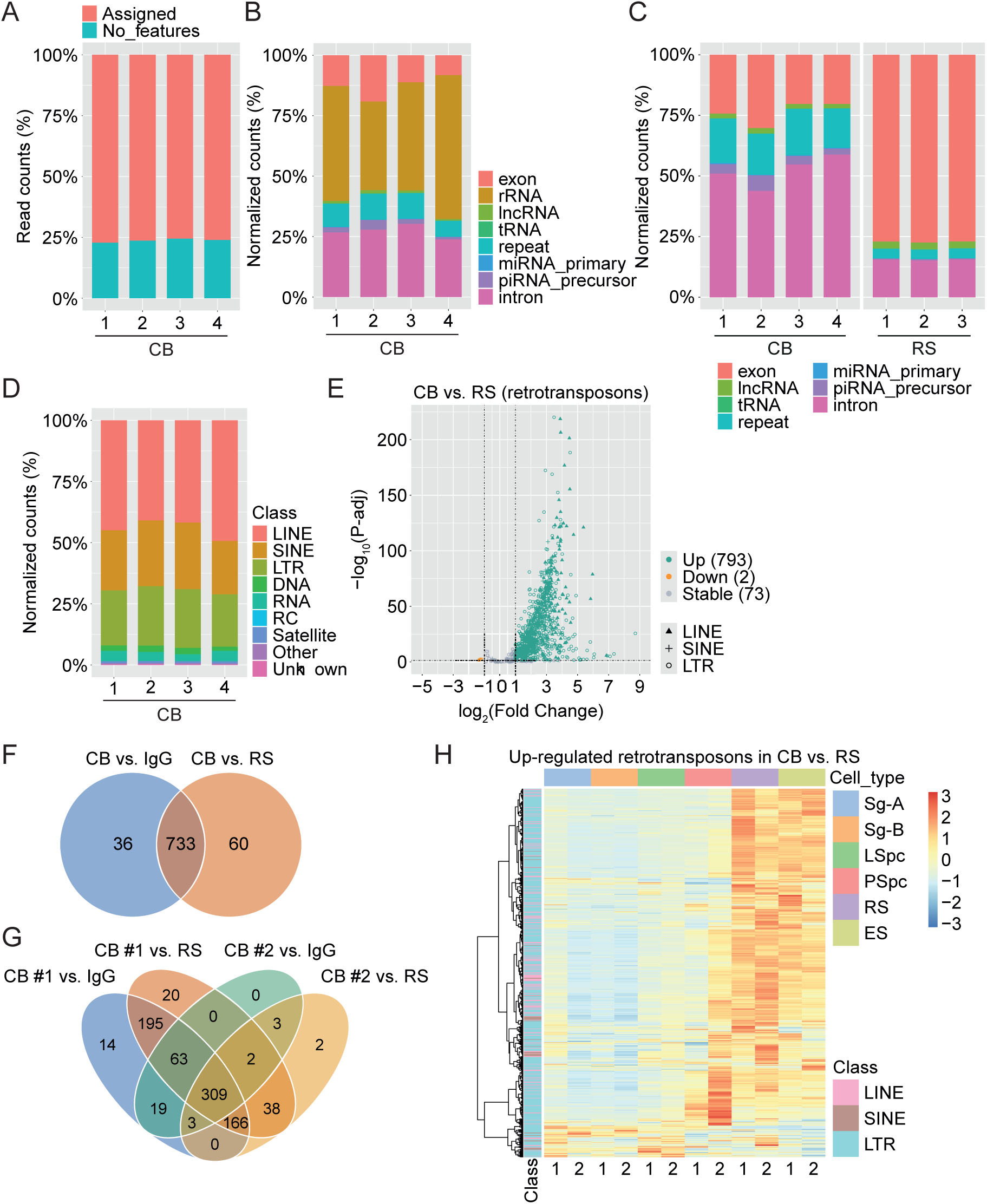
The CB enriches transposable elements. (A) Distribution of RNA-seq reads between the annotated (Assigned) and non-annotated (No_features) parts of the genome in CBs (batch 1) isolated from adult mouse testes. (B) Read distribution across RNA biotypes in adult CBs (batch 1). (C) Read distribution across RNA biotypes (excluding rRNA reads) in adult CBs (batch 1) and round spermatids (RS). (D) Read distribution across different classes of repeats in the CBs (batch 1) from adult mouse testes. (E) Volcano plot of differentially expressed retrotransposons between adult CB (batch 1) and RS (P-adj ≤ 0.05, |log₂FC| ≥ 1). (F) Venn diagram showing the overlap between retrotransposons enriched in the CB (adult, batch 1) vs. IgG controls and the CB (adult, batch 1) vs. RS. (G) Venn diagrams showing the overlap of up-regulated retrotransposons identified in two independent batches of adult CBs. Comparisons include CB (batch 1) vs. RS, CB (batch 1) vs. IgG, CB (batch 2) vs. RS and CB (batch 2) vs. IgG. A total of 309 mRNAs are shared among all four comparisons and defined as high-confidence CB-enriched retrotransposons. (H) Heatmap showing expression dynamics of the CB-enriched retrotransposons (batch 1, CB vs. RS) during spermatogenesis (GSE35005). Rows represent genes and columns represent spermatogenic stages. Sg-A: type A spermatogonia, Sg-B: type B spermatogonia, LSpc: leptotene spermatocytes, PSpc: pachytene spermatocytes, RS: round spermatids, ES: elongating spermatids.

### Transposable element transcripts are highly enriched in the CB

The relative higher proportion of repeat-derived reads in the CB compared to the input round spermatid samples (Fig. 2C) was intriguing, considering the central role of the CB in the piRNA pathway, with well-documented role in transposon silencing in the fetal germline ^18^. This prompted us to further explore the accumulation of repeat transcripts in the CB. We showed that the majority of the CB-associated repeat-derived reads mapped to retrotransposons (LINE, SINE and LTR families) (Fig. 2D), and we therefore focused the following differential expression analysis on this class of repeats. The analysis identified prominent enrichment of retrotransposons in the CB compared to round spermatids (793 out of 868) (Fig. 2E). When compared to IgG controls, 769 out of 878 retrotransposons were enriched in the CB, with 733 shared between the two comparisons (Fig. 2F). Replication using an independent batch of CBs isolated from adult testes at different time by a different researcher (n = 2) ^19^ identified 317 retrotransposons enriched in CBs compared to both round spermatids and IgG controls, of which 309 overlapped with those detected in the first dataset (Fig. 2G), validating the repeatability of the finding. Developmental expression analysis (GSE35005) revealed that most of the CB-enriched retrotransposons (from Fig. 3E) had an expression peak in spermatids (Fig. 2H), suggesting that the CB primarily accumulates repeat-derived RNAs expressed during the haploid phase, rather than residual transcripts from earlier phases.

**Figure 3.**
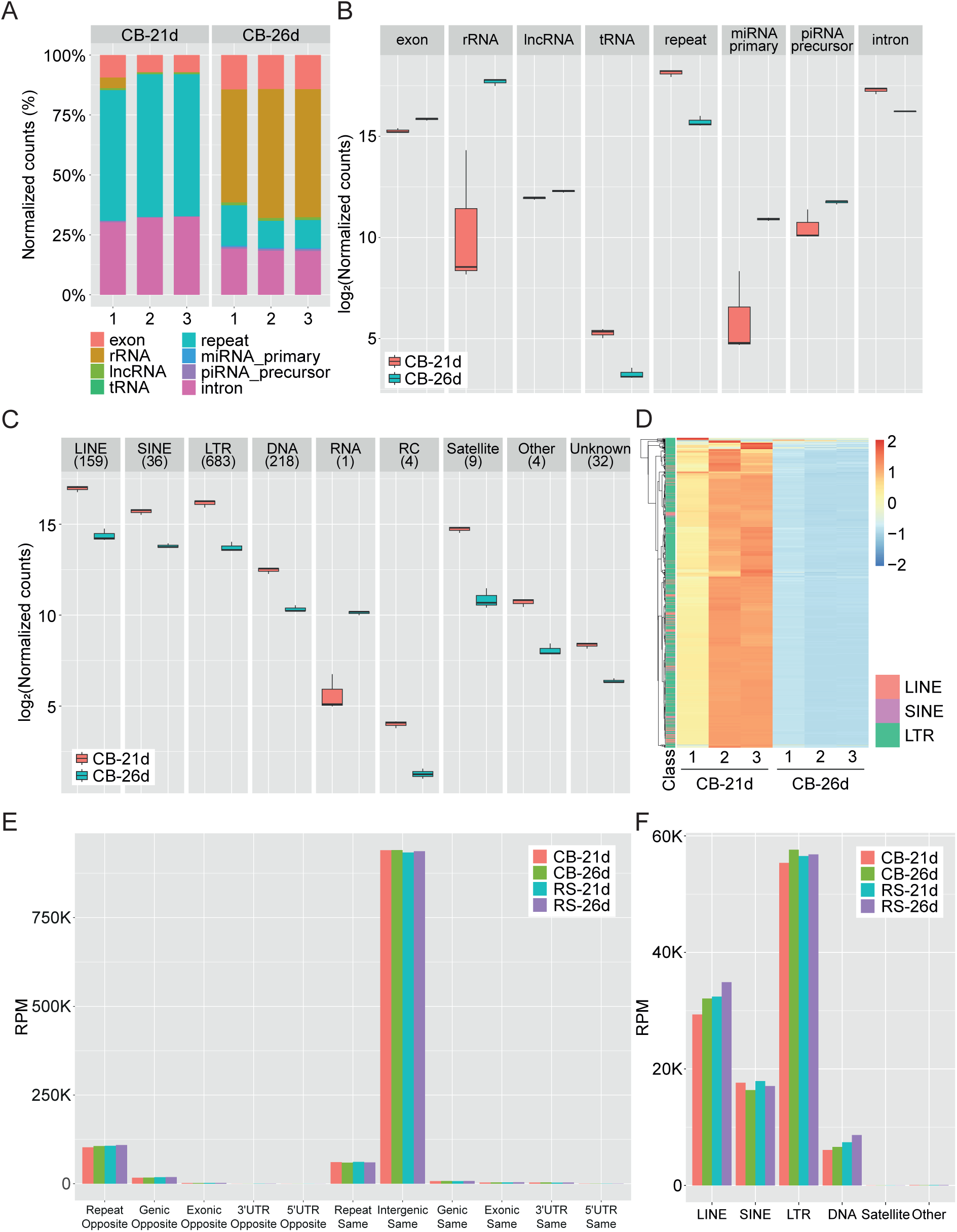
The RNA composition of the CB is remodeled during round spermatid differentiation. (A) Read distribution across RNA biotypes in CB-21d and CB-26d. (B) Number of normalized reads across RNA biotypes in CB-21d and CB-26d. (C) Abundance of different classes of repeats in CB-21d and CB-26d. (D) Heatmap showing expression differences of retrotransposons between CB-26d and CB-21d. (E) Origins of pachytene piRNA reads classified by genomic annotation of CB-21d, CB-26d, RS-21d and RS-26d. (F) Origins of antisense repeat-derived pachytene piRNA reads classified by repeat classes of CB-21d, CB-26d, RS-21d and RS-26d.

### Transposable element transcripts are more prominently enriched in early CBs

To understand the differentiation-associated changes in the CB RNA composition, we performed RNA-seq for the CBs isolated from early-(21 dpp) and late-phase (26 dpp) round spermatids and respective negative control IgG IP samples (Supplementary Fig. S4A). Principal component analysis showed distinct clustering of samples from 21 and 26 dpp groups, suggesting substantial differences in their transcriptomic profiles (Supplementary Fig. S4B). Read distribution analyses revealed intriguing differences between the 21 dpp and 26 dpp CBs, with the early-CB containing a higher proportion of reads from repetitive elements, while the proportion of the rRNA-originated reads was higher in the 26 dpp CB (Fig. 3A). Comparison of the normalized read counts from different RNA subtypes verified the increase in rRNA, miRNA precursor and piRNA precursor reads and decrease in tRNA, repeat and intronic reads in the CB when progressing from early to late round spermatid phase (Fig. 3B). These findings indicate dynamic changes in the CB RNA composition as round spermatid differentiation progresses.

The presence of relatively more repeat-derived reads in the early 21 dpp CB compared to 26 dpp CB motivated us to assess how CB-associated repeat transcript profile changes during spermatid differentiation. Quantification of normalized read counts originating from different repeat classes showed that the 21 dpp CB contained more reads from all classes of retrotransposons (LINE, SINE, LTR) (Fig. 3C). Differential expression analysis revealed that the majority of retrotransposons were significantly enriched in both 21 dpp CB and 26 dpp CBs (862 and 815, respectively) compared to IgG controls (Supplementary Fig. S4C,D). However, the direct comparison of 26 dpp and 21 dpp CBs demonstrated that almost all retrotransposons (851) were significantly more abundant in 21 dpp CBs (Fig. 3D). The observed decrease in repeat abundance of retrotransposons transcripts in 26 dpp CBs may reflect either enhanced processing efficiency or reduced transcription of repeat loci as round spermatid differentiation proceeds.

Interestingly, mapping of the CB-localized mature piRNAs from the same time points (21 dpp and 26 dpp) to the genome revealed that, while most reads originated from the sense strand of intergenic regions, typical for pachytene piRNAs, a smaller fraction mapped to repetitive elements in both sense and antisense strands (Fig. 3E). We next focused on piRNAs mapping to the antisense strand of repetitive elements and assessed their expression across repeat classes. The resulting expression profiles showed that most of these piRNAs are antisense to retrotransposons (Fig. 3F). These piRNAs may contribute to the regulation of transposable element transcripts enriched within CBs by directing sequence-specific silencing mechanisms. Collectively, these findings support a role for the CB in suppressing potentially deleterious transposable element expression during spermatid development.

### CB-enriched mRNAs are functionally linked to RNA regulation and meiotic processes

Next, we focused on protein-coding gene-derived exonic reads to identify mRNAs specifically enriched in the CB. We used two different approaches for the gene-level differential expression analysis to compare the CB either to the input round spermatid samples or to the negative IgG control, and identified 4161 and 4333 significantly enriched genes, respectively (Fig. 4A, Supplementary Fig. S5A, Supplementary Table S2A,B). 2395 of these genes were found to be enriched in the CB in both comparisons and were defined as CB-enriched genes (Fig. 4B, Supplementary Table S2A,B). These genes exhibited stage-dependent expression patterns when examined across the spermatogenic trajectory (GSE35005) ^20^ (Supplementary Fig. S5B). Majority of them were relatively less expressed in haploid cells compared to earlier cell types (expression groups 1 and 2). Group 1 genes were expressed during all phases of spermatogenesis until pachytene spermatocyte phase, and group 2 expression peaked in pachytene spermatocytes (Supplementary Fig. S5B). The downregulation of their expression coinciding with the CB-localization of the gene products suggests that they may be expressed in meiotic cells and translocated to the CB for degradation after meiotic divisions. The remaining one-fourth peaked in round and elongating spermatids, indicating that they may be targeted to the CB right after their synthesis in haploid cells, for example for temporal translational regulation, a process known to be active at this phase of differentiation ^1^. GO enrichment analysis of CB-enriched genes revealed strong enrichment for functions related to RNA splicing and RNA regulation, as well as chromosome segregation and other processes related to meiotic division (Supplementary Fig. S5C, Supplementary Table S2C).

**Figure 4.**
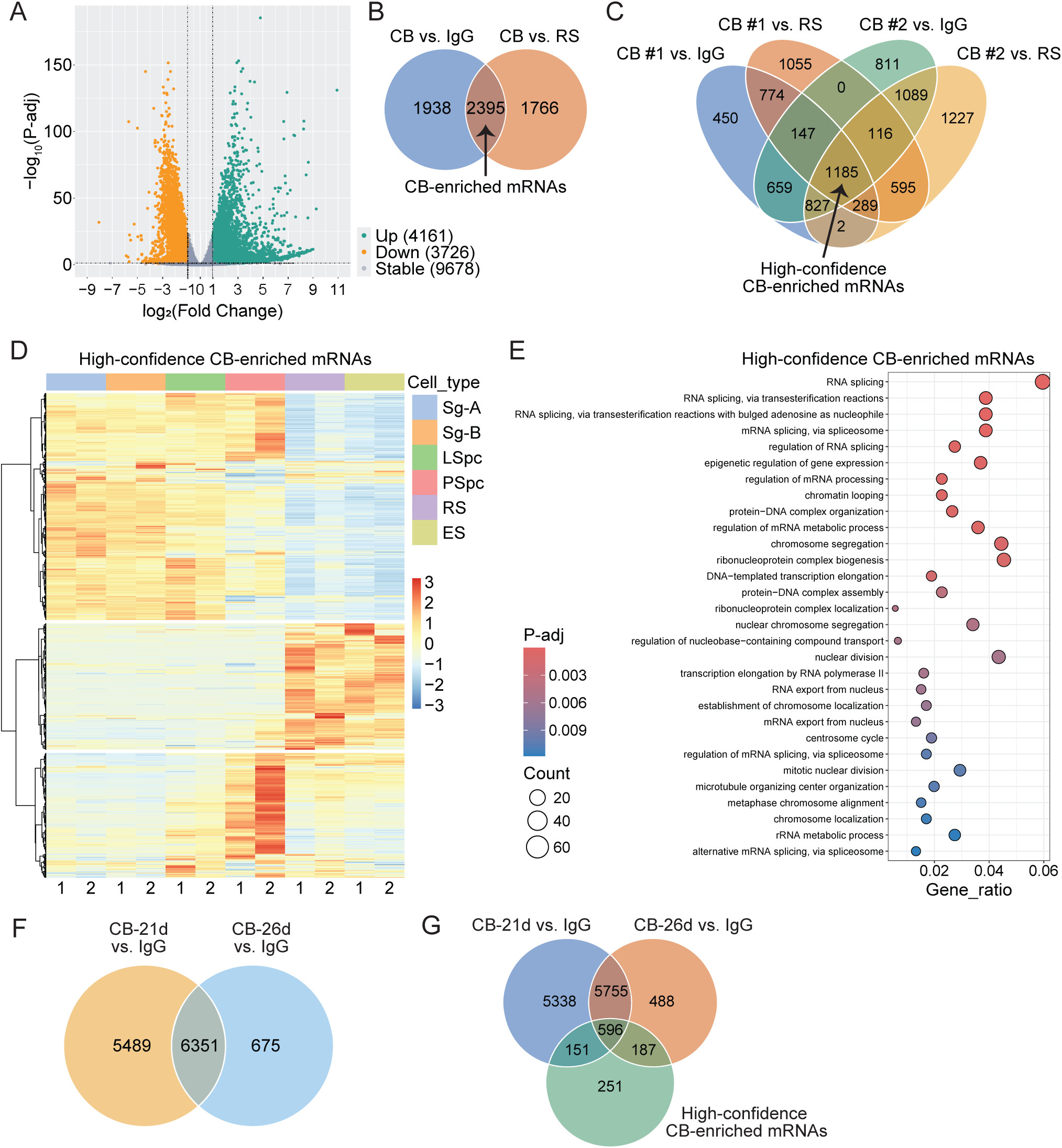
The CB enriches a distinct set of mRNAs and the CB composition is dynamically modified during differentiation. (A) Volcano plot of differentially expressed mRNAs between adult CB (batch 1) and round spermatids (RS) (P-adj ≤ 0.05, |log₂FC| ≥ 1). (B) Venn diagram showing overlap of up-regulated mRNAs identified in adult CB (batch 1) vs. RS and adult CB (batch 1) vs. IgG. (C) Venn diagram showing the overlap of up-regulated mRNAs identified in two independent CB isolation experiments. Comparisons include adult CB (batch 1) vs. RS, adult CB (batch 1) vs. IgG, adult CB (batch 2) vs. RS and adult CB (batch 2) vs. IgG. A total of 1185 mRNAs are shared among all four comparisons and defined as high-confidence CB-enriched genes. (D) Heatmap showing the relative expression levels of high-confidence CB-enriched genes across spermatogenesis (GSE35005). Rows represent genes and columns represent spermatogenic stages. Sg-A: type A spermatogonia, Sg-B: type B spermatogonia, LSpc: leptotene spermatocytes, PSpc: pachytene spermatocytes, RS: round spermatids, ES: elongating spermatids. (E) GO enrichment analysis of biological process terms for high-confidence CB-enriched mRNAs. Top 30 enriched terms are shown. (F) Venn diagram showing the overlap of up-regulated mRNAs between CB-21d vs. IgG and CB-26d vs. IgG. A total of 6351 mRNAs are shared between both stages, while 5489 and 675 mRNAs are uniquely enriched in CB-21d and CB-26d, respectively. (G) Venn diagram showing overlap among CB-21d vs. IgG, CB-26d vs. IgG, and adult high-confidence CB-enriched mRNAs. A total of 596 mRNAs is shared across all three datasets.

To assess the robustness of these findings, we used again an independent batch of adult CBs (n = 2) ^19^ for the same analysis pipeline. This replication identified 3217 CB-enriched genes (shared between CB vs. round spermatid and CB vs. IgG comparisons), 1185 of which overlapped with the first dataset and were defined as high-confidence CB-enriched mRNAs (Fig. 4C, Supplementary Table S2A,B). These mRNAs displayed similar spermatogenic expression patterns (Fig. 4D) and functional enrichments (Fig. 4E) as the CB-enriched mRNAs from the first sample set (Supplementary Fig. S5B,C). The reproducibility across biological replicates demonstrates that the CB consistently associates with mRNAs produced from the specific subset of genes in round spermatids.

### Dynamic remodeling of CB transcriptome accompanies round spermatid differentiation

To clarify the dynamics of the CB mRNA cargo during round spermatid differentiation, we then focused on exonic reads from protein-coding genes at distinct steps of round spermatid differentiation, and identified 11840 and 7026 genes significantly enriched in 21 dpp CB and 26 dpp CB compared to IgG controls, respectively (Supplementary Table S3A,B). Of these, 6351 were shared between both CB stages, suggesting a conserved RNA core, while 5489 and 675 mRNAs were specifically enriched in 21 dpp CB and 26 dpp CB, respectively (Fig. 4F). Therefore, despite the slightly lower total exonic read abundance (Fig. 3B), the CB in early round spermatids appears to accumulate a much more diverse repertoire of mRNAs than the CB at later steps of differentiation. GO enrichment analysis revealed that mRNAs specifically enriched in the CB at early (21 dpp) vs. later (26 dpp) steps of round spermatid differentiation represented very distinct biological processes (Supplementary Fig. S6A,B, Supplementary Table S3C,D), reflecting the functional shift of the CB during differentiation. Integration with the adult CB datasets showed that the majority of the high-confidence adult CB-enriched mRNAs were identified also in 21 dpp CBs (747) and 26 dpp CBs (783), with 596 shared across all three datasets (Fig. 4G). This conserved set likely represents core mRNA persistently associated with the CB throughout spermatogenesis. All together, these results indicate that the CB is not a static RNA repository but a dynamically remodeled RNP granule that selectively recruits specific mRNA populations, and the majority of mRNAs recruited to the CB in early round spermatids are later removed from the CB either by degradation or release to the cytoplasm.

### Intron-retaining transcripts accumulate in the CB after meiotic division

The enrichment of intronic reads in the CB (Fig. 2C) suggests that the CB may associate with incompletely spliced pre-mRNAs. To further study the intron retention in the CB, we performed a global alternative splicing analysis comparing the adult CB transcriptome to round spermatid input. The change in Percent Spliced In (ΔPSI) was used to quantify the differential splicing events. The analysis revealed a striking global shift toward intron retention in the CB. Specifically, we identified 3254 significantly intron retention events (FDR ≤ 0.05 and |ΔPSI| ≥ 0.1) (Supplementary Table S4A). The majority of the events (3174/3254, 97.5%) showed increased intron retention in the CB (Fig. 5A). The 3174 CB-enriched intron retention events mapped to 1817 genes, of which 320 overlapped with the previously identified CB-enriched genes from Fig. 4B. Gene Ontology (GO) enrichment analysis of these 320 genes showed a strong enrichment of genes encoding for splicing regulators (Supplementary Fig. S6C, Supplementary Table S4B). The expression of most of them was shown to peak in late meiotic cells (Supplementary Fig. S6D), which is in line with the reported high intron retention activity in meiotically expressed transcripts ^21^. Comparison of the CBs from early (21 dpp) and late (26 dpp) phases of round spermatid differentiation showed that the intronic reads were highly represented in the CB at both time points (Fig. 3A,B). However, 21 dpp CBs were shown to contain somewhat more intronic reads than 26 dpp CBs (Fig. 3A,B). These results suggest that intron-containing transcripts produced in meiotic cells accumulate in the CB after meiotic division, and they are retained in the CB and degraded/released from the CB later during differentiation.

**Figure 5.**
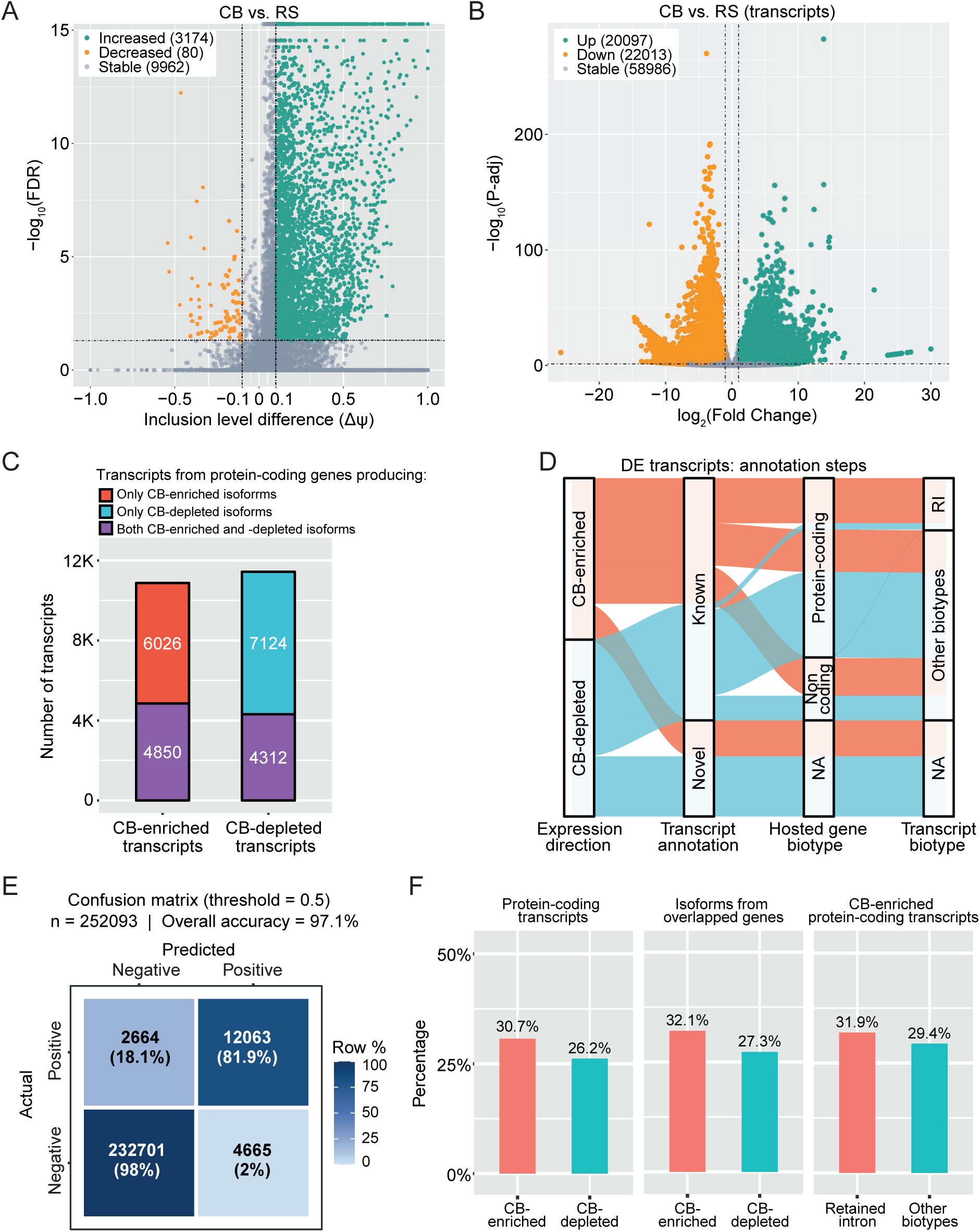
CB enriches isoforms with retained introns. (A) Volcano plot of intron retention events between adult CB (batch 1) and round spermatids (RS) (FDR ≤ 0.05, |inclusion level difference| ≥ 0.1). (B) Volcano plot Volcano plot showing differential transcript expression between CB (batch 1) and RS based on assembled transcripts (P-adj ≤ 0.05, |log₂FC| ≥ 1). (C) Stacked bar plot showing the gene-level origins of CB-enriched and CB-depleted transcripts. A substantial proportion of transcripts originate from genes producing both enriched and depleted isoforms (purple), indicative of isoform switching. (D) Sankey diagram tracking the flow of differentially expressed transcripts (CB-enriched and CB-depleted) through successive annotation layers: transcript annotation (known vs. novel), hosted gene biotype (protein-coding, non-coding and NA) and transcript biotype (retained intron (RI), other biotypes and NA). NA: not available. (E) Confusion matrix evaluating the performance of the piX-Plore deep learning model on a held-out mouse piRNA-mRNA test set (n = 252093) at a threshold of 0.5, demonstrating 97.1% overall accuracy. (F) Bar charts showing the percentage of transcripts predicted to be targeted by piRNAs (binding score ≥ 0.99) across various comparisons: all DE transcripts produced from protein-coding genes (left), DE transcripts produced from overlapping genes (middle), and CB-enriched transcripts produced from protein-coding genes and annotated as retained intron (right). Targeting rates remain consistently near ∼30% across all transcript subgroups.

### Specific isoforms of genes are selectively enriched in the CB

The enrichment of intron-retaining transcripts in the CB prompted us to investigate whether the CB selectively accumulates specific isoforms from individual genes. To address this question, we assembled transcripts from the adult CB and round spermatid RNA-seq datasets (GSE182518) and performed transcript-level differential expression analysis. This analysis identified 20097 transcripts that were significantly upregulated in the CB relative to the round spermatid input, hereafter referred to as CB-enriched transcripts (Fig. 5B, Supplementary Table S5). Conversely, 22013 transcripts were significantly downregulated and were classified as CB-depleted transcripts (Fig. 5B, Supplementary Table S5).

Approximately 25% of the CB-enriched transcripts represented previously uncharacterized, unannotated transcripts, whereas the remainder corresponded to known isoforms, including 10876 transcripts produced from annotated protein-coding genes. Notably, a more detailed analysis of protein-coding genes revealed widespread isoform-specific enrichment within the CB. Specifically, for 2737 genes, certain isoforms were enriched in the CB (4850 isoforms), while alternative isoforms from the same genes were depleted (4312 isoforms) (Fig. 5C, Supplementary Table S5). Interestingly, supporting our earlier finding on the upregulation of intron retention events in the high-confidence CB-enriched genes (Fig. 5A), we showed that almost 60% (2799 out of 4850) of these CB-enriched isoforms were annotated as “retained intron” (Fig. 5D), compared to only 7% of “retained intron” isoforms among the CB-depleted transcripts. These findings demonstrate that, in addition to the selective post-transcriptional regulation of particular genes in the CB (*e.g.* high-confidence CB-enriched genes), specific isoforms from the same gene can be differentially targeted to or excluded from the CB, with intron-retention serving as a prominent CB-targeting signal.

### piRNA-targeting potential does not explain CB-enrichment

To study if the CB-enriched pachytene piRNAs could be involved in recruiting longer RNAs into the CB by sequence-specific targeting, we developed an *in silico* tool piX-Plore, a novel deep learning framework, to predict piRNA-targeting sequences. Model performance was evaluated on a held-out test set of mouse piRNA-transcript pairs. At a standard binding score threshold of 0.5, the model demonstrated an overall accuracy of 97.1%, precision of 72.1% and recall of 81.9% (Fig. 5E, Supplementary Fig. 7A-C). To ensure maximum biological confidence and strictly minimize false positives, we applied a highly stringent binding score threshold of ≥ 0.99, retaining only the highest-confidence interactions for downstream analysis.

We selected piRNAs for the targeting analysis from the adult CBs by mapping them into the experimentally defined piRNA clusters ^15^. Furthermore, we specifically selected only pachytene piRNAs due to their dominance in the CB over the pre-pachytene piRNAs. Then we used a threshold of 10 reads to select only the most abundant pachytene piRNAs in the CB, and only the piRNAs that were expressed in all replicates were kept for downstream analysis. In total of 172986 piRNAs were left after the filtering, and they mapped to 100 different pachytene piRNA clusters.

Next we predicted the potential of these selected piRNAs to target CB-enriched (10876) vs. CB-depleted (11436) protein-coding transcripts. Analysis of the distribution of these highly confident piRNA-mRNA interactions (binding score threshold of ≥ 0.99) across different transcript populations showed that one third of the CB-enriched transcripts were targeted by piRNAs (30.7%). However, only slightly reduced proportion (26.2%) of CB-depleted transcripts were identified as piRNA targets (Fig. 5F), therefore, piRNA-targeting potential cannot differentiate these two populations. We then narrowed down the analysis to those genes producing both CB-enriched and CB-depleted isoforms, this baseline targeting rate remained consistent (32.1% vs. 27.3%, Fig. 5F). Furthermore, the presence of retained introns in the transcript did not alter targeting frequency, as CB-enriched transcripts annotated as “intron retained” showed nearly identical targeting rate (31.9%, Fig. 5F). Together, these consistent proportions suggest that piRNAs target a broad, basal fraction of the transcriptome (∼30%), rather than acting as the primary driver for the specific enrichment or depletion of mRNAs in the CB.

### CB-localization of transcripts is encoded in sequence features

To determine whether transcript localization to the CB is encoded by intrinsic sequence properties, we asked whether sequence-derived features alone could distinguish CB-enriched from CB-depleted transcripts. Using a defined population of 20097 CB-enriched and 22013 CB-depleted transcripts (Fig. 5B), we evaluated whether any sequence-derived features could accurately predict transcript destination, including at the level of individual isoforms originating from the same genetic locus. Interestingly, an XGBoost classifier trained on the full set of 862 sequence-derived features accurately distinguished CB-enriched from CB-depleted transcripts, achieving a mean 5-fold cross-validated ROC AUC of 0.8419 (Fig. 6A). Therefore, transcript destination can be predicted with high accuracy from sequence features alone, indicating that CB localization is largely encoded within the RNA itself.

**Figure 6.**
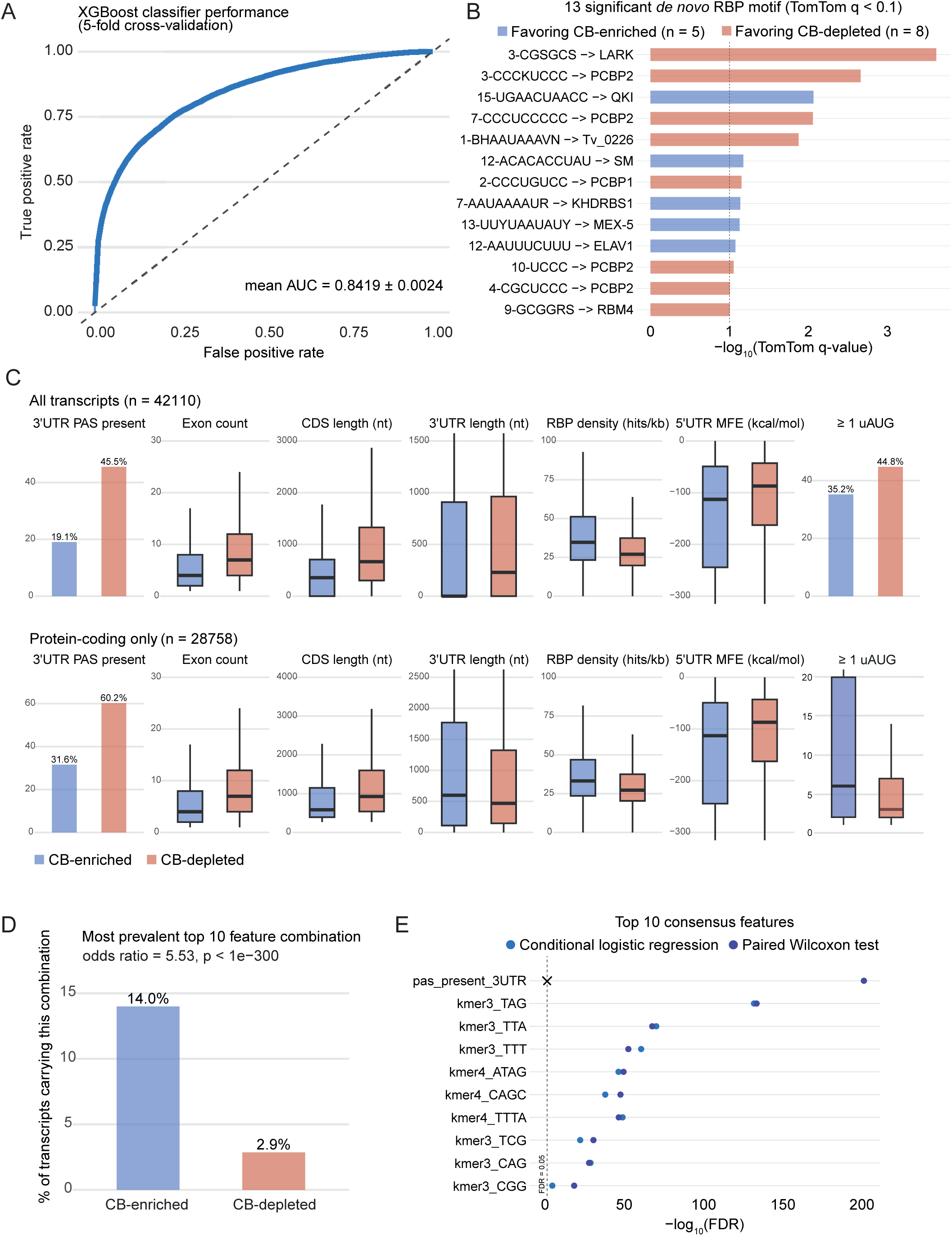
CB-enriched transcripts are distinguished by specific sequence features. (A) XGBoost classifier performance. Receiver Operating Characteristic (ROC) curve evaluating the performance of an XGBoost classifier using 5-fold cross-validation. The model achieves a mean Area Under the Curve (AUC) of 0.84. (B) Significant *de novo* motifs stratified by enrichment direction. Bar chart showing the 13 *de novo* motifs identified by STREME that matched RNA-binding protein motifs in the Ray2013 RNAcompete database (TomTom q < 0.1). Motifs are ranked by -log10(TomTom q-value). Blue bars indicate motifs prevalent in CB-enriched transcripts, and red bars indicate motifs prevalent in CB-depleted transcripts. (C) Comparison of key sequence and structural features between CB-enriched (blue) and CB-depleted (red) transcripts. The top row shows analysis across all assembled transcripts (unfiltered, n = 42110), while the bottom row focuses strictly on protein-coding transcripts possessing an Open Reading Frame (has_orf = TRUE, n = 28758). Features compared include the percentage of transcripts containing a 3’UTR Polyadenylation Signal (PAS), total exon count, Coding Sequence (CDS) length, 3’UTR length, RNA-Binding Protein (RBP) binding site density, and 5’UTR Minimum Free Energy (MFE). For the final panel (≥ 1 uAUG), the top row shows the percentage of transcripts carrying at least one upstream AUG (uAUG) in the 5′UTR; in the bottom row, the distribution of uAUG count among only the transcripts that carry ≥1 uAUG. (D) Bar chart illustrating the prevalence of the top 10 most predictive feature combinations within CB-enriched vs. CB-depleted transcripts. This specific architectural combination is highly enriched in CB transcripts (14.0% vs. 2.9%; odds ratio = 5.53, P-adj < 1e-300). (E) Top 10 consensus features remain significant after controlling gene of origin. Dot plot showing whether each of the top 10 features still statistically distinguishes CB-enriched from CB-depleted transcripts when the comparison is made strictly within the same gene using two independent statistical tests. Nine out of ten features remained significant by both tests (dashed line represents the significance cutoff); the tenth PAS could not be evaluated by one of the two tests but remained significant by the other.

K-mer composition analysis revealed distinct nucleotide signatures between the two groups. 575 of 640 tested k-mer features (89.8%) were significantly differentially represented after FDR correction (Supplementary Fig. S8A). CB-depleted transcripts were characterized by GC-rich k-mers (*e.g*., CCA, ACC, ACG), whereas CB-enriched transcripts showed marked enrichment for AU-rich k-mers, including poly-U tracts and TAG-context motifs (*e.g*., TAG, TTA, TTTA). Independent *de novo* motif discovery using STREME (Supplementary Fig. S8B) recapitulated these patterns without relying on a predefined k-mer vocabulary, consistently identifying AU-rich motifs among CB-enriched transcripts and GC/CG-rich motifs among CB-depleted transcripts across all four region-orientation analyses. Notably, the canonical AAUAAA polyadenylation signal (consensus BHAAUAAAVN) was significantly enriched among CB-depleted transcripts (11193 sites; E = 1.7 × 10^-69^). Of 32 motif/RBP pairings with a reportable TomTom comparison, 13 reached statistical significance (q < 0.1, Fig. 6B). Among these was HuR/ELAVL1, an AU-rich element-binding protein whose cognate motifs were enriched in CB-associated transcripts. This observation is particularly noteworthy because HuR has previously been reported to localize to the CB and to mediate trafficking of AU-rich mRNAs during spermiogenesis, providing an independent biological link between the identified sequence signatures and CB function ^7,22^. Collectively, these findings indicate that CB-enriched and CB-depleted transcripts possess distinct sequence architectures and identify AU-rich regulatory elements as prominent features of CB-associated RNAs.

### Compartment identity is defined by a combinatorial feature set

Although sequence composition strongly predicted transcript destination, no individual feature was sufficient to explain compartment identity on its own. To determine how different transcript properties contribute to localization, we interrogated the model using SHAP analysis. This analysis revealed that CB localization is specified by a combinatorial architecture of sequence and structural features rather than by a single dominant determinant (Fig. 6C). The strongest individual contributor was the presence of a polyadenylation signal (PAS) within the terminal 100 nt of the 3′UTR. Such a signal was present in 45.5% of CB-depleted transcripts (10021/22013) but only 19.1% of CB-enriched transcripts (3830/20097) (odds ratio 3.55, 95% CI 3.40-3.71; P-adj < 0.01) (Fig. 6C).

Beyond PAS usage, CB-enriched transcripts exhibited a distinct structural profile. They contained fewer exons (median 4 vs. 7) and shorter coding sequences (median 357 vs. 666 nt; P-adj < 0.05) than CB-depleted transcripts (Fig. 6C). They also displayed a higher density of predicted RBP-binding motifs (median 34.7 vs. 27.0 hits/kb; P-adj < 1 × 10^-5^), more stable folding of their 5′UTRs (median MFE -112.9 vs. -87.0 kcal/mol; P-adj = 2.9 × 10^-10^), and a greater number of upstream AUG codons among transcripts containing at least one uAUG (median 6 vs. 3; P-adj = 7.7 × 10^-28^). When all transcripts were considered together, CB-enriched transcripts appeared to possess shorter 3′UTRs. However, CB-enriched transcripts were also substantially more likely to lack a predicted ORF (39.7% vs. 24.4%), confounding this comparison. Restricting the analysis to ORF-containing transcripts reversed this relationship: CB-enriched transcripts contained significantly longer 3′UTRs than CB-depleted transcripts (median 599 vs. 468 nt; P-adj = 7.6 × 10^-9^) (Fig. 6C), suggesting that expanded 3′UTR sequence space may provide additional opportunities for localization-related regulatory interactions.

Finally, we asked whether these features act independently or in concert. Consistent with a combinatorial model, the consensus features accumulated rather than substituted for one another. The most common co-occurrence pattern among the median-binarized top ten features was itself strongly associated with CB localization, occurring in 14.0% of CB-enriched transcripts (2814/20097) compared with only 2.9% of CB-depleted transcripts (630/22013) (Fig. 6D). Notably, this effect exceeded that observed for any individual feature alone. These results indicate that CB localization is governed by a multivariate sequence code in which multiple transcript properties act cooperatively to bias compartmental destination.

### The sequence code operates at the isoform level

A key question is whether the identified sequence features reflect an intrinsic transcript-level sorting code or merely gene-level differences in expression and regulation. To distinguish between these possibilities, we focused on genes that produced both CB-enriched and CB-depleted transcript isoforms. Among the 18356 genes represented in our dataset, 2737 (14.9%) generated transcripts assigned to both compartments, contributing 9363 transcripts and yielding 8135 within-gene CB-enriched/depleted transcript pairs (Supplementary Fig. S8C,D). This design enabled direct comparison of alternative isoforms derived from the same locus, effectively controlling for gene identity and shared transcriptional environment.

Remarkably, nine of the ten consensus localization features remained significantly associated with transcript destination after controlling for gene of origin (FDR < 0.05, Fig. 6E). The PAS feature could not be tested by conditional logistic regression because of quasi-complete separation within small, matched gene-strata. However, it remained significant by paired-Wilcoxon test alone (FDR < 0.01). Thus, the same sequence characteristics that distinguish transcripts globally also discriminate alternative isoforms generated from a common gene. To assess the practical magnitute of this within gene discrimination, we used the full sequence feature classifier to ask how often the CB-enriched transcript of a matched pair scored higher than its CB-depleted sibling. The classifier correctly ranked 81.4 % of the CB-enriched/CB-depleted pairs. These findings demonstrate that CB localization is specified at the level of individual isoforms rather than at the level of gene identity. Taken together, our results support a model, in which compartment destination in round spermatids is encoded by a combinatorial and graded sequence code that operates directly on individual RNA molecules.

## DISCUSSION

The CB has long been recognized as a defining feature of post-meiotic male germ cells and has been implicated in diverse RNA regulatory processes during spermatogenesis ^6,9,10,23^. Although its protein composition has been extensively characterized, the molecular principles governing RNA recruitment to the CB, as well as the function of this specialized germ granule, have remained poorly understood. In this study, we combined transcriptomic and small RNA profiling of isolated CBs across distinct stages of round spermatid differentiation to define the RNA landscape of the CB and uncover mechanisms underlying its RNA recruitment. Our findings provide a comprehensive framework for understanding how the CB organizes distinct RNA populations and reveal previously unappreciated changes in its molecular composition during spermatogenesis.

A major conclusion of this study is that the CB functions as an active RNA-sorting and regulatory compartment rather than a passive repository of cytoplasmic transcripts. Its enrichment for piRNA pathway-associated RNAs, transposable element-derived transcripts and specific mRNA isoforms demonstrates a striking degree of transcript selectivity. Furthermore, the extensive reorganization of CB RNA content during spermatid differentiation suggests that CB-mediated regulatory processes are continuously adapted to developmental stage-specific requirements. Together, these findings identify the CB as a developmentally regulated center of RNA metabolism that coordinates multiple post-transcriptional pathways in haploid male germ cells.

One prominent feature of the CB transcriptome was the strong enrichment of transposable element transcripts, supporting a role for the CB in transposon control during haploid differentiation. CB-associated transposon transcripts predominantly originated from LINE and LTR families and displayed maximal expression during haploid differentiation. Importantly, transposon transcripts were particularly abundant in early CBs, indicating that their sequestration is most prominent immediately after meiosis. The enrichment of transposon transcripts provides a compelling functional link between the CB and the piRNA pathway that has a well-established function in transposon silencing in the fetal germline ^18,24^. Consistent with previous studies ^7,9^, we observed strong accumulation of PIWI proteins and pachytene piRNAs within the CB, and showed that a fraction of the CB-associated piRNAs are complementary to transposon sequences. Importantly, our analyses further revealed that the CB-associated pachytene piRNA population remains remarkably unchanged throughout round spermatid differentiation, forming a relatively stable framework upon which developmental regulation is superimposed.

The enrichment of transposon transcripts in the CB raises the question of why this regulation may be particularly important during post-meiotic differentiation. Spermatogenesis involves extensive chromatin remodeling, culminating in the histone-to-protamine transition and dramatic reorganization of the genome ^25^. Such transitions may increase the risk of transposon activation and genomic instability ^26,27^. By concentrating both transposon transcripts and PIWI–piRNA complexes within the same compartment, the CB may function as a specialized surveillance center that preserves genome integrity when conventional epigenetic silencing mechanisms are challenged.

Beyond transposon regulation, our study revealed extensive association of the CB with protein-coding mRNAs. This observation is particularly intriguing given previous reports demonstrating that pachytene piRNAs can regulate protein-coding transcripts ^28–31^. The CB function has also been linked to the mRNA regulation through the NMD pathway ^9,32^. We identified a reproducible core set of CB-enriched genes across independent datasets, indicating that recruitment of specific mRNAs to the CB is not stochastic but represents a conserved feature of spermatid differentiation. The dynamic behavior of these mRNAs provides important insight into CB function. Early round spermatid CBs contained a substantially broader mRNA repertoire than late CBs, whereas a subset of transcripts persisted throughout differentiation. Combined with previous evidence demonstrating the localization of the NMD factor SMG6 to the CB and its reported role in mRNA regulation during the transition from meiotic to post-meiotic gene expression programs ^9^, these observations support a model in which many transcripts are initially targeted to the CB shortly after meiosis for surveillance and degradation, while a smaller subset escapes decay and remains associated with the compartment during later stages of differentiation. Thus, the CB may function as a critical regulatory platform where transcripts undergo surveillance and are directed toward degradation, storage, or eventual reuse.

Our transcript-level analyses further demonstrate that CB targeting is fundamentally isoform-specific. Thousands of genes generated both CB-enriched and CB-depleted transcript variants, indicating that compartmentalization is not determined at the level of gene identity. Instead, distinct isoforms produced from a common locus can be differentially sorted between the CB and the surrounding cytoplasm. This finding considerably expands the regulatory capacity of the CB and reveals an additional layer of post-transcriptional control operating during spermatid differentiation.

Interestingly, we showed that CB-enriched transcripts were highly enriched for retained introns, suggesting that intron retention may function as an important determinant of CB recruitment or retention. Regulated intron retention has previously been described in meiotic germ cells, where many intron-containing transcripts are unusually stable and can later associate with ribosomes during spermiogenesis ^21^. Our data therefore support a model in which intron-retaining transcripts are selectively routed to the CB following meiosis. Whether CB localization facilitates their eventual maturation and utilization or instead commits them to degradation remains unresolved. Notably, genes encoding splicing regulators were particularly enriched among CB-associated intron-retaining transcripts. This observation raises the possibility that the CB participates in autoregulatory feedback loops that modulate splicing capacity during the transition from meiotic to post-meiotic development. Given the absence of canonical spliceosomal machinery from the CB ^7^, elucidating the ultimate fate of these intron-retaining transcripts will be an important direction for future studies.

Pachytene piRNAs and PIWI proteins are highly enriched within the CB, and previous studies have demonstrated that piRNA-loading is important for localization of PIWIL1 to the CB ^13^ and pachytene piRNAs can direct mRNA regulation in spermatids ^29,31^. These observations raise the possibility that piRNA-target interactions could contribute to transcript recruitment into the CB. However, the comparable predicted piRNA-targeting potential of CB-enriched and CB-depleted transcripts argues that piRNA recognition alone is insufficient to explain CB localization. Notably, in *Drosophila* germ plasm, Aub-bound piRNAs were shown to act as an "adhesive trap" that tethers mRNAs to germ granules through partial base pairing ^33^. While our findings suggest that a similar mechanism is unlikely to drive transcript recruitment to the mammalian CB, it remains possible that CB-associated piRNAs contribute to the retention of localized RNAs and facilitate their subsequent surveillance, storage or degradation.

While piRNA-target recognition did not explain transcript recruitment to the CB, our findings point instead to transcript-intrinsic determinants as the primary drivers of localization. Specifically, CB localization was encoded by a combination of AU-rich sequence motifs, RNA structural characteristics, and 3′UTR-associated features rather than by any single determinant. The existence of such a sequence code suggests that intracellular RNA organization in spermatids can be specified directly by transcript architecture, providing an additional layer of post-transcriptional regulation beyond gene expression itself. Such a mechanism would allow closely related RNA molecules to be differentially handled without altering overall transcriptional programs.

In summary, our findings establish the CB as a highly selective post-transcriptional regulatory compartment that coordinates the handling of distinct RNA populations during spermatogenesis. By recognizing intrinsic features of individual transcripts, the CB provides a mechanism for transcriptome remodeling through selective RNA partitioning and processing. The functional significance of this regulatory activity is reflected in numerous mouse models, where disruption of CB integrity or associated pathways leads to impaired spermatid maturation and male infertility ^5,6^. These observations underscore the importance of CB-mediated RNA regulation for reproductive competence and suggest that CB dysfunction may contribute to human reproductive disorders.

## MATERIALS AND METHODS

### Mice

Mice from C57BL/6 strain were used in the experiments. Mice were maintained and housed at the Central Animal Laboratory of the University of Turku and the Finnish Animal Ethics Committee approved all experiments.

### Isolation of CBs

21 dpp, 26 dpp and adult mice were sacrificed, testes were collected, and CBs were isolated using the previously published protocol ^7,9^. Briefly, germ cells were released from testes by incubation in 0.05% (w/v) collagenase (Worthington). The cell suspension was passed through a 100-µm strainer (BD Falcon), washed with phosphate-buffered saline (PBS), and cross-linked in 0.1% paraformaldehyde (Electron Microscopy Sciences) for 20 min at RT. Fixed cells were lysed by sonication (UCD-200, Diagenode) in 1 ml RIPA buffer (50 mM Tris-HCl, pH 7.5; 1% NP-40; 0.5% sodium deoxycholate; 0.05% SDS; 1 mM EDTA; 150 mM NaCl; supplemented with protease inhibitors (Roche), 0.2 mM PMSF, and 1 mM DTT) using six 30 sec pulses at medium power. The CB-enriched fraction was collected by centrifugation at 600 × g for 10 min, resuspended in RIPA buffer, and sonicated for two additional 30 sec intervals. Lysates were combined and then split equally for the CB immunoprecipitation with anti-DDX4 antibody (Abcam, ab13840) and the negative control immunoprecipitation with rabbit IgG (Cell Signaling Technology, 2729S), using Protein G Dynabeads (Thermo Fisher Scientific, cat. no. 10003D). A small fraction of the sample was used for the Western blotting, and the majority was used for RNA extraction. For the isolation of 21 dpp and 26 dpp CBs, in total 18 mice were sacrificed per time point. 9 mice were used for CB isolation for RNA-seq and 9 for small RNA-seq. The material from three mice were pooled into one sample (n = 3 biological replicates per group). For the adult CB small RNA-seq, one sample contained material from one mouse (n = 3 biological replicates).

### Isolation of Intermitochondrial cement (IMC)

For IMC isolation, the protocol described by Olotu et al. (2023) was used ^12^. Testes from six 15-day-old male mice were collected in PBS, and seminiferous tubules were digested in 0.5 mg/ml collagen type I (LS004196, Worthington Biochemical) in PBS containing 0.1% glucose and 1 µg/ml DNase I (LS006353, Lakewood, NJ, USA) for 60 min at room temperature with rotation. Tubules were filtered through a 100 µm strainer (BD Biosciences), washed with 0.1% glucose in PBS, and cross-linked with 0.1% (v/v) paraformaldehyde (Thermo Fisher Scientific) for 20 min at room temperature. Cells were centrifuged at 300 × g for 5 min at 4°C and lysed by sonication (Diagenode UCD-200) in RIPA buffer (50 mM Tris-HCl pH 7.5, 1% NP-40, 0.5% sodium deoxycholate, 0.05% SDS, 1 mM EDTA, 150 mM NaCl, protease inhibitors, 0.2 mM PMSF, 1 mM DTT). Lysates were centrifuged at 300 × g for 10 min at 4°C, pre-cleared with Dynabeads Protein G (Invitrogen), and subjected to immunoprecipitation overnight at 4°C with 2 µg antibody or IgG control. Bead-bound complexes were washed three times with lysis buffer and used for small RNA-seq and Western blotting.

### Isolation of round spermatids

Round spermatids were isolated using the previously published protocol ^34^. For each biological replicate, testes were collected from six 21 dpp and six 26 dpp mice. Decapsulated testes were first digested with collagenase IV (Sigma, C5138), followed by a second digestion with trypsin (Worthington, LS003703) and DNase I (Sigma-Aldrich, DN25) in 1× KREBS buffer. The resulting cell suspension was washed with 1× KREBS, filtered through a 100 µm cell strainer, and loaded onto a pre-chilled BSA gradient. Following sedimentation, germ cell fractions were collected, washed, and their purity was assessed by DAPI staining. Round spermatid fractions 1-12 (65-80% purity) were collected, cells isolated from three mice were pooled into one sample (n= two biological replicates per time point), and RNA was extracted and sent for small RNA-seq.

### Western blotting

Protein samples were denatured in Laemmli buffer at 95°C for 5 min and separated by SDS-PAGE on 4–20% Mini-PROTEAN TGX precast gels (Bio-Rad, cat. no. 4561093). Proteins were transferred to PVDF membranes (Amersham, cat. no. RPN303F) by wet transfer. Membranes were blocked with 5% skimmed milk in 0.1% TBST for 1 h at room temperature and incubated overnight at 4°C with primary antibodies diluted in the blocking solution. After washing with 0.1% TBST, membranes were incubated with HRP-conjugated anti-rabbit or anti-mouse IgG (1:1000) in the same buffer for 1 h at room temperature. Signals were detected using Western Lightning ECL Pro (PerkinElmer, cat. no. NEL120E001EA), imaged with a LAS4000 system (FujiFilm), and quantified using ImageJ (NIH, version 1.8.0).

### Immunofluorescence

WT-like adult mouse testes were fixed in 4% phosphate buffered formaldehyde overnight. The fixed samples were dehydrated, embedded into paraffin and cut to sections. Paraffin-embedded testis sections were deparaffinized by incubating 3 × 5 min in xylene, 2 × 10 min in 100% ethanol, 2 × 10 min in 96% ethanol, 2 × 10 min in 70% ethanol and washed 2 × 2 min in milliQ water. Sections were subjected to antigen retrieval in Tris–EDTA buffer (10 mM Tris base, 1 mM EDTA Solution, 0.05% Tween 20, pH 9.0) by boiling in pressure cooker at 120°C for 20 min. Sections were then blocked with 10% bovine serum albumin (Sigma, A2153) in PBST (0.1% Tween-20) for 1 h. Primary antibodies against MIWI (Cell Signaling, #2079) and MILI (Cell Signaling, #2071) were labeled with FlexAble CoraLite® Plus 488 and 594 Antibody Labeling Kits for Rabbit IgG (Proteintech, KFA001 and KFA009), respectively. 0.5 μg of both antibodies were labeled according to the manufacturer’s protocol, resulting in the volume of 10 μl. Both labeled primary antibodies (1:100 dilution in blocking solution) and an anti-DDX4 antibody (R&D Systems, AF2030, 1:200 dilution in blocking solution) were incubated together with tissue sections overnight at +4 °C in a humidified environment. Alexa Fluor donkey anti-goat 647 secondary antibody (Thermo Fisher Scientific, A-21447, 1:500 dilution in blocking solution) was incubated for 1 h at room temperature in a humidified environment. All washes were done with 0.1% PBST. Sections were mounted with ProLong™ Diamond Antifade Mountant (Thermo Fisher Scientific, P36961). Z-stack images were obtained with 3i CSU-W1 Spinning disk confocal microscope with 100x objective. Acquired images were deconvolved with Huygens Professional software (version 25.10.1p1, 64-bit, Scientific Volume Imaging, Hilversum, The Netherlands) using automatic deconvolution parameters and a theoretical point spread function, and further processed using ImageJ (version 1.54r, National Institute of Health, USA) and Adobe Photoshop.

### RNA extraction and sequencing

RNA was extracted from enriched fractions of round spermatids or isolated CBs with the TRIzol reagent (Thermo Fisher Scientific) using standard protocols. Isolated RNA was analyzed using a NanoDrop (Thermo Scientific) and Bioanalyzer (Agilent). The DNase I (Sigma-Aldrich, AMPD1) was used to remove genomic DNA from the samples. Non-stranded total RNA libraries were prepared from CBs isolated at 21 dpp (n = 3) and 26 dpp (n = 3), as well as IgG IP control samples from 21 dpp (n = 3) and 26 dpp (n = 3). Quantified libraries were sequenced by NovaSeq X Plus Series with 150 bp pair-end at Novogene (UK) Company Limited. For small RNA-seq, CBs from adult (n = 3), 21 dpp (n = 3) and 26 dpp (n = 3) testes, and round spermatids from 21 dpp (n = 2) and 26 dpp (n = 2) testes were subjected to small RNA library preparation and sequenced on the NovaSeq 6000 platform (1 × 50 bp).

### Gene-level RNA-seq analysis

The reads quality were first evaluated by FastQC (v0.11.9) (https://www.bioinformatics.babraham.ac.uk/projects/fastqc/). The adapters were trimmed off from raw reads using trimmomatic (v0.39) ^35^. Then the clean reads were mapped to the mouse reference genome (Ensembl: GRCm38) using STAR (v2.7.10a) ^36^. The reads were assigned and counted using featureCounts (v2.0.3) ^37^. The normalization and differentially expression analysis were done by DESeq2 (v1.48.0) package in R (v4.5.2) ^38^. The genes with P-adjust value (P-adj) ≤ 0.05 and |log₂FC| ≥ 1 were considered as the differentially expressed genes. The reads were also assigned to several sets of sequences for read distribution by custom annotation which includes protein-coding genes, long non-coding RNAs from GENCODE ^39^, tRNA fragments from UCSC, rRNAs from UCSC RepeatMask, miRNA primaries from miRbase ^40^, piRNA precursors from piRBase ^41^, repeats (https://labshare.cshl.edu/shares/mhammelllab/www-data/TEtranscripts/TE_GTF/) and introns. Alternative splicing events were analyzed by rMATS (v4.3.0) ^42^. The magnitude of splicing change between groups was expressed as the Delta PSI (ΔPSI, or Δψ), representing the difference in mean inclusion levels between two groups (ψA−ψ). To ensure robust quantification, events were filtered to require a minimum of 10 total supporting junction reads (the sum of inclusion and skipping counts across all replicates) per event. TEtranscripts and TEcounts were used for analyzing repeats including transposable elements ^43^.

### Transcript-level RNA-seq analysis

The transcripts were assembled for each CB and round spermatid sample from adult mice separately by StringTie (v3.0.3) with mouse genome annotation (Ensembl: GRCm38) ^44^. The annotations were merged and reads were counted based on the merged annotation. The transcripts with total counts more than 10 across all samples were kept. DESeq2 was used for transcript normalization and differential expression analysis ^38^.

#### Data processing and feature extraction

Transcripts assembled with StringTie were extracted using gffread against the GRCm38 mouse genome. Coding sequences (CDS) and 5′/3′ untranslated region (UTR) boundaries were predicted using TransDecoder. Following quality filtering, the final dataset comprised 42110 transcripts (20097 CB-enriched and 22013 CB-depleted). We computed 862 features per transcript to characterize sequence and structural properties. These included: transcript, 5′UTR, CDS, 3′UTR lengths, exon count, open reading frame (ORF) status, 5′UTR upstream-AUG (uAUG) abundance, polyadenylation hexamer (AAUAAA/AUUAAA) presence and position relative to the 3′ end, and 3-mer/4-mer nucleotide composition (transcript-wide and 3′UTR-specific). Composition features were computed by exhaustively enumerating all 4^k^ possible k-mers (k = 3, 4) over the nucleotide alphabet, tallying occurrences with a single-base sliding window, and normalizing by the number of valid k-mer positions per sequence to obtain a length-independent frequency. Motifs in the 5′ and 3′ UTRs were identified *de novo* using STREME (MEME suite v5.5.9), comparing CB-enriched sequences against CB-depleted sequences as background and vice versa, and matched against the Ray2013 RNAcompete database (244 motifs) using TomTom. The complete Ray2013 library, together with the 60 *de novo* motifs discovered, was scanned genome-wide against all transcripts using FIMO (P-adj < 1 × 10^-4^); resulting hit counts were length-normalized to hits per kilobase. Minimum free energy (MFE) was calculated for 5′ and 3′ UTRs (capped at 3000 nt for computational tractability) using RNAfold (ViennaRNA) and included as a predictive feature, together with its length-normalized form.

#### Bias controls and model validation

To mitigate transcript-length bias, RNA-binding protein (RBP) and *de novo* motif hit counts were length-normalized, reducing their correlation with transcript length from r = 0.83 to r = 0.003. To exclude a sequencing-depth artifact, we confirmed that including DESeq2-derived transcript abundance (baseMean) as a classifier feature inflated ROC AUC from 0.842 to 0.904; baseMean was excluded from all models reported below, isolating classification performance attributable strictly to sequence-intrinsic features. We trained an XGBoost classifier (v3.1.2; 300 trees, maximum depth 5, learning rate 0.05, subsample 0.8, colsample_bytree 0.5) to discriminate CB-enriched from CB-depleted transcripts, evaluated by stratified 5-fold cross-validation (scikit-learn v1.8.0). Feature importance was assessed by mean absolute SHAP value (shap v0.52.0) on a 5000-transcript subsample. As an independent line of evidence, features were also ranked by LASSO logistic regression (L1-regularized, regularization strength selected by internal cross-validation) and random forest importance, with predictive performance assessed by repeated (5×5-fold) stratified cross-validation.

### Small RNA-seq analysis

The small RNA-seq analysis pipeline was modified from the previous study ^9^. The Cutadapt (v4.1) was used to remove the adapter and low quality of the raw reads ^45^. The size of clean reads was subsequently filtered and only reads between 23 and 35 nt in length were retained. These filtered reads were mapped to the mouse genome (GRCm38) by HISAT2 (v2.2.1) ^46^. The reads mapped to tRNA and rRNA locations were removed from the BAM files. The featureCousnts (v2.0.3) was used for assigning the remaining reads to piRNA cluster annotations ^37,47^. The reads that were successfully mapped to the piRNA cluster regions were considered as the piRNAs. In addition to the individual piRNAs, we also did piRNA cluster analysis based on the piRNA reads. The normalization and differential expression analysis of individual piRNAs or piRNA clusters were done by DESeq2. The SPORTS (v1.1) was only used to investigate the read distributions by mapping the clean reads to miRNA (miRBase), tRNA derived small RNAs (tsRNAs) (GtRNAdb), rRNA derived small RNAs (rsRNAs) (rRNAdb) and piRNA (piRBase) ^48^. The SPORTS results were not used for any other following analysis.

### piRNA target prediction

Potential piRNA-mRNA interactions were predicted using piX-Plore (manuscript in preparation), a deep learning framework for genome-scale piRNA target prediction. piX-Plore employs a windowed transformer architecture that partitions full-length transcripts into overlapping 2000 nucleotide windows with a 500-nucleotide stride, scoring each window independently and assigning the maximum window-level score as the transcript level binding probability. To ensure robust performance in mammalian systems, the model was initialized using weights pre-trained on *Caenorhabditis elegans* experimental piRNA interaction data generated using CLASH-seq and 5′ RACE-based approaches ^49,50^ and subsequently fine-tuned on *Mus musculus* data compiled from published studies ^29–31,51^ and from piRBase ^52^. Negative pairs were generated by cross-pairing approximately 18265 unique MIWI-pathway piRNAs with transcriptionally neutral transcripts identified from GSE70731 ^53^. Model performance was evaluated on a held-out test set of mouse piRNA-transcript pairs not seen during training. The model achieved a window-level ROC-AUC of 0.984 (Supplementary Fig. S7A), with an average precision of 0.857 (Supplementary Fig. S7B). At a binding score threshold of 0.5, the model demonstrated an overall accuracy of 97.1%, precision of 72.1%, recall of 81.9%, F1 score of 0.767, and Matthews Correlation Coefficient of 0.753 (Supplementary Fig. S7C). Full benchmarking, architectural details, and a web application will be described in the accompanying tool manuscript. Interactions with a binding score ≥ 0.99 were retained for downstream analysis. Source code is available at: https://github.com/simang5c/piX-plorer.

## DATA AVAILABILITY

RNA-seq data produced in this study are deposited in NCBI BioProject database under accession number PRJNA1493897.

## ACKNOWLEDGEMENTS

We thank all Kotaja lab members for their support and help. We thank Matthieu Bourgery for providing the foundational pipeline used for small RNA-seq data processing, which was modified for this study. The authors wish to acknowledge CSC - IT Center for Science, Finland, for computational resources. Turku Central Animal Facility is acknowledged for providing facilities for animal maintenance and experimentation.

## AUTHOR CONTRIBUTION

N.K. and L.M. conceived and designed the project. A.A. collected the samples and performed the experiments. L.M. and S.C. performed bioinformatic analysis. S.L. and O.O. performed the experiments. N.K. supervised the study. N.K. and L.M. wrote the paper. All authors contributed to finalizing the manuscript.

